# Between specialization and flexibility: how tasks and space shape behavioral profiles in ant colonies

**DOI:** 10.64898/2026.07.19.739471

**Authors:** Pol Fernández-López, Jolle W. Jolles, Daniel Oro, Frederic Bartumeus

## Abstract

Task allocation in eusocial insects has long been studied under the framework of division of labor, implying a relatively rigid association between individuals and tasks. However, most eusocial species lack morphological specialization, and workers regularly switch tasks as colony demands change. This raises a fundamental question: do tasks shape the behavioral profiles of workers, or does individual behavioral variation cut across task boundaries? We addressed this in a controlled laboratory study of *Aphaenogaster senilis* ants, comparing the behavioral profiles of four task groups (scouts, recruits, nurses, and necrophores) spatially segregated by their location within the colony setup and subsequently tested individually in four ecologically relevant contexts. This multivariate profiling, still rarely applied in ants, revealed that some tasks impose clear behavioral specialization (scouting, brood care), whereas others do not (recruitment, necrophoresis). Critically, this specialization appears in foraging-related tasks, whereas sociality does not: it varies considerably among workers, even within a single task group. Behavioral specialization, therefore, exists, but not across every dimension of behavior, and it is not a fixed property of the task. These results suggest that workers may differ in their readiness to shift roles depending on the task at hand, and this variation may in turn shape how colonies adapt to environmental change. More broadly, our results speak to a question central to collective behavior research well beyond ants: how individual variability translates into functional structure at the group level.

## 1 Introduction

Eusocial insects function as decentralized cognitive systems in which information processing emerges from local interactions among workers [1, 2, 3]. As individuals move and interact, the structure of these interactions dynamically reshapes, forming a fluid and adaptive information network, a ‘liquid brain’ [4, 5, 6]. Through simple behavioral rules and encounters that vary with local density, colonies coordinate large-scale patterns such as resource allocation, activity rhythms, and task performance at the collective level [7, 8, 9]. A central question is whether such fluid self-organization is supported by highly structured, task-specialized behavioral profiles or whether it instead relies on flexible behavioral repertoires that allow individuals to adapt rapidly to changing colony needs.

The most studied route to such organization is division of labor, the repartition of tasks among workers, which is shaped by genetic background, life history, physiology, and age polyethism [10, 11]. In some taxa, division of labor is reinforced by morphological adaptations, usually linked to body size (e.g., major and minor workers in *Atta* spp. or *Pheidole* spp., [12, 13]), such that body form largely predetermines the tasks a worker performs. Yet, morphological cases are the exception: more than 70% of ants are monomorphic [14], and their workers assemble into functionally distinct tasks that reorganize as colony demands change [15, 16, 17, 18]. In these species, then, division of labor must arise without morphological structuring, raising the question of what distinguishes workers engaged in different tasks, if at all.

One candidate mechanism is individual behavioral heterogeneity, which shapes a wide range of ecological, evolutionary, and social processes across solitary, social, and eusocial species [19, 20, 21]. In eusocial insects, such heterogeneity can manifest across multiple organizational scales, from individual workers [22, 23, 24] to functional groups [25, 26, 27], and whole colonies [28, 29]. This multi-scale structure is consistent with the superorganism perspective in which the colony is the primary unit of function, evolution, and fitness [17]. However, whether this heterogeneity is organized by task remains unresolved: workers performing the same task may share characteristic behavioral profiles or individuals may differ more from one another than from the task groups they belong to.

Addressing this has been difficult because it requires tracking many individual ants across multiple tasks, an approach still rarely applied in ants (but see [25, 26, 27]). Here we address this gap in *Aphaenogaster senilis* ants, an omnivorous, generalist, monomorphic species whose foragers exhibit remarkable behavioral heterogeneity [30, 31, 22, 6, 32]. To assign workers to task groups, we drew on the concept of activity spaces, originally developed in the human sciences to describe the set of locations an individual regularly visits, each associated with specific activities [33, 34]. An analogous principle applies to eusocial insects, where tasks are tied to specific regions of the nest and its surroundings, generating predictable spatial–task associations. Exploiting this, we spatially segregated four task groups - scouts (autonomous foraging), necrophores (corpse disposal), recruits (cue-guided foraging), and nurses (brood care) - and characterized each worker’s behavioral profile across four ecologically relevant assays: two foraging-related (exploration and responsiveness to a liquid food source), and two communication-based (sociability and responsiveness to chemical cues). Beyond testing each behavior in isolation, we combined these assays with a multi-dimensional analysis of behavior [35, 36], mapping each worker’s full behavioral profile into a common behavioral space to ask whether task identity structures how individuals behave across ecological contexts. Although well established in other fields like cell biology [37], such behavioral mapping has rarely been applied to social insects, where it is well suited to resolving whether behavior is organized by task. Resolving whether behavior is structured by task in a monomorphic species helps clarify how eusocial colonies remain both robust, through consistent task performance, and flexible, through the capacity to reallocate workers when conditions change.

## 2 Materials and Methods

### 2.1 Subjects and housing

In spring 2025, we collected four queenright *Aphaenogaster senilis* colonies from a suburban, partially vegetated area near Cerdanyola, Spain. Each colony contained between 300 and 800 workers, along with brood at various developmental stages, including eggs, larvae, and pupae. Two colonies were used for piloting to ensure the assays elicited the desired responses and produced sufficient behavioral heterogeneity, while the remaining two were kept naive and used for the experiments. All colonies were transferred to our climate-controlled laboratory, maintained under a 14:10 light–dark cycle at (25 ±2°C), and housed in custom-designed nest spaces compartmentalized by function (Fig. 1).

**Figure 1:**
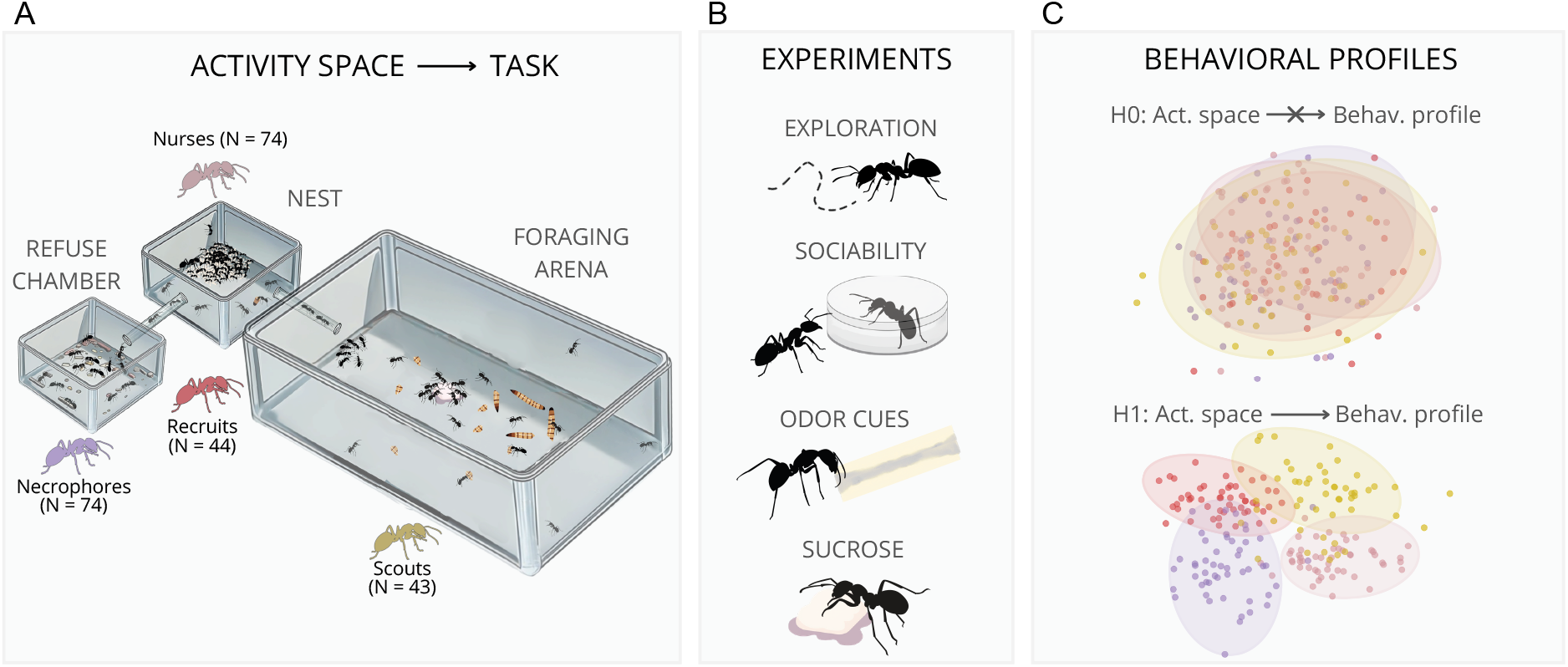
Main rationale of our work, from the mapping of activity spaces onto tasks, and tasks onto behavioral profiles. (A) Each activity space is expected to allocate a main task: necrophores in the refuse chamber, nurses in the nest, recruits transitioning from the nest to the foraging arena (within and in the vicinity of the tube), and scouts in the foraging arena. (B) We collected ants from each of the locations represented in (A) and individually tested them across four conditions: exploratory activity, degree of sociability, responsiveness to chemical cues, and responsiveness to a liquid food source. From each of these experiments, several metrics were used to build a multivariate behavioral profile that reflects each individual’s propensity to explore, engage in (direct or indirect) social interactions, and feed. (C) Conceptualization of the two contrasting predicted outcomes from the main hypothesis tested in the present work. If the tasks carried out by workers do not predict their behavioral profiles (null hypothesis, H0), the behavioral profiles should be distributed over a shared behavioral space. Otherwise (alternative hypothesis, H1), the distinct behavioral profiles should form clusters in the behavioral space, showing minimal overlap among the activity spaces.

Each nest comprised a main chamber connected to a refuse chamber, and could be temporarily joined to an open foraging arena via a connecting tube (Fig. 1). The main chamber (69 × 69 × 24, mm) housed the brood, queen, and the nurse workers responsible for brood care. It was covered with an opaque cloth, and water was provided *ad libitum* (replenished weekly) to maintain a humid, dark environment. The adjoining refuse chamber (65 × 45 × 25, mm) served as a hygienic zone for waste segregation and necrophoric behavior, and was perforated and exposed to light to approximate conditions outside the nest. To promote necrophoresis, we placed some dead individuals and debris in this chamber. Three times per week, the main chamber was connected for approximately one hour each morning, via the connecting tube, to an open foraging arena (430 × 280 × 70mm) with Fluon-coated walls to prevent ant escape. This temporary connection stimulated foraging, with scouts expected to roam the arena and recruits to remain near the nest or within the connecting tube. Each feeding session provided dehydrated *Tenebrio molitor* larvae as a protein source, supplemented once a week with apple pieces or sucrose solution for carbohydrates.

### 2.2 Worker classification

We assigned workers to four task groups (scouts, recruits, nurses, and necrophores) based on the activity space in which they were located at the time of collection. This approach rests on the well-documented spatial fidelity of eusocial insect workers: individuals tend to occupy consistent regions of the nest and its surroundings over time, with their spatial position closely tied to the tasks they perform (e.g., [38, 39, 40, 41, 26, 42, 43, 44, 45]). The worker’s location provides a practical, non-invasive proxy for task identity, yet we acknowledge that spatial zones are unlikely to map perfectly onto tasks and each location probably contains a mixture of task types in differing proportions.

One hour prior to the experiments, the colony was connected to the foraging arena to allow foragers to leave the nest naturally, simulating a routine feeding session. Following this period, workers from each activity space were carefully isolated, with precautions taken to minimize colony disturbance and prevent stress or undesired movement between compartments. First, the foraging arena was disconnected from the main nest to isolate and collect recruits, defined as individuals located within the connecting tube or clustered within 5 cm of the entrance of the feeding arena. The remaining workers within the foraging arena were then collected and classified as scouts. Next, the refuse chamber was disconnected to collect the necrophores. Finally, nurses were carefully collected from the main nest, prioritizing individuals actively engaged in brood-carrying behavior.

Ants were collected from two colonies over four experimental days, with 60 individuals collected per day (120 per colony total), targeting approximately 15 workers per behavioral group (scouts, recruits, nurses, necrophores). Because scouts and recruits leave the nest voluntarily, we could not consistently reach 15 individuals per group for these two categories; the daily quota of 60 was therefore completed by collecting additional nurses and necrophores. Across the four days, this yielded 240 ants in total, of which 5 were excluded from analysis due to death or loss during handling, leaving a final sample of 235 workers (43 scouts, 44 recruits, 74 nurses, 74 necrophores). After collection, each worker was transferred to an individually labeled Eppendorf tube, and its compartment of origin was recorded. Ants were left to acclimate in the tube for 10 minutes before undergoing the first behavioral assay (exploration). An additional 30 nurses were collected each day, separately from the focal sample, to serve as social stimuli in the sociability assay. Nurses were chosen for this role because they represented the most numerous group in the colony, and using a single behavioral category standardized the social stimuli across setups, avoiding a mixed composition of nurses, necrophores, and other groups within the same social chamber. Three nurses were placed in the social chamber of each of the ten experimental setups.

### 2.3 Experimental setup

The assays were conducted using an array of ten experimental boxes, each containing a white acrylic testing arena (75 × 55 × 25 cm). These boxes provided visual and acoustic isolation, ensuring tested individuals were undisturbed for the duration of each trial. Arenas were uniformly illuminated from above with diffused LED lighting, and the walls were coated with Fluon to prevent escape. Each arena was filmed from above using a Raspberry Pi 3B+ and associated v2 noir HD camera (Raspberry Pi Foundation, Cambridge, UK) above a diffuser panel, and all arenas were thoroughly cleaned with 70% ethanol at the end of each experimental day. We designed four behavioral assays to quantify individual responses across key ecologically and functionally relevant scenarios: a novel open environment, conspecifics, chemical cues, and liquid food (Fig. 1).

#### Exploration assay

To measure exploratory behavior and the response to a novel environment, the focal ant was tested in the open arena without any further cues provided.

#### Sociability assay

To characterize social attraction and propensity for social interactions, a group of three untested nurses was placed inside a social chamber inside the testing tanks. The social chamber consisted of a small, upside-down Petri dish (diameter 3.5 cm) perforated to allow the transfer of visual and tactile cues. Three individuals provided a sufficient social stimulus while minimizing stress on the ants and avoiding the difficulty of trapping larger groups. Nurses were used to standardize the stimulus, as their brood-carrying behavior is reliable and easily identified. These nurses used as social stimuli in this essay were separate individuals collected for this purpose and were not among the tested workers. At the start of each trial, the focal ant was placed immediately adjacent to the social chamber to ensure immediate exposure to the social stimulus.

#### Chemical cue assay

To assess responsiveness to chemical communication, which ants commonly use to coordinate colony behavior [46], we deposited a gaster-extract cue trail in the arena. The extract was prepared by homogenizing three ant gasters in 2 mL of methanol (following [47]). Although its precise composition was not determined, it is expected to contain a cocktail of trail and pheromones as described in [48], and our pilots confirmed it reliably elicited clear behavioral reactions in *A. senilis*. Our aim was to characterize the behavioral responses triggered by the cue, and to test whether a common stimulus of similar concentration and composition elicited different reactions among task groups. A strip of filter paper (10 cm × 2 cm) was coated with a line of the extract and was placed partially inside the acclimation Petri dish so that the focal ant could detect the cue before being released. The strip was replaced after every trial to maintain a consistent concentration across individuals.

#### Sucrose assay

To quantify feeding motivation and persistence as a proxy for foraging behavior and energy demand, at the start of the trial, the focal ant was placed in the acclimation Petri dish together with a piece of cotton soaked in 1M sucrose solution, which was then subsequently removed. As *Aphaenogaster senilis* does not share liquid food through trophallaxis, this assay was designed to assess the metabolic state (degree of hunger) of the tested individual and the perceived value of the food source.

### 2.4 Experimental procedure

Experiments were run over four consecutive days in daily batches of 60 trials (Colony 1 on Days 1–2; Colony 2 on Days 3–4). To reduce the presence of unwanted chemical cues, assays were conducted in a strict sequential order based on stimulus intensity. Exploration and sociability assays were performed first and second, respectively, when the arena floor was free of cues except for potential residual cuticular hydrocarbons. These were followed by chemical cue (third) and sucrose (fourth) assays; neither the pheromone nor the sucrose was ever directly deposited on the arena floor, thereby preserving data integrity throughout the experimental workflow.

The pipeline was structured by assay type rather than by testing individual ants through all assays consecutively. Within each daily batch, assays were conducted in sequential blocks: the exploration assay was run for all 60 individuals first, followed in turn by the sociability, chemical cue, and sucrose assays. For each trial, a focal ant was selected from its Eppendorf tube in a pre-defined randomized order and introduced into one of the ten arenas, where it was initially confined under an inverted Petri dish (3.5 cm diameter), placed in the center for the exploration assay, or in a randomized location otherwise. After one minute of acclimation, the Petri dish was carefully lifted, and a 10-minute video-recording started at 10 fps using standardized, optimized settings with PiRecorder [49]. The ant was then returned to its labeled Eppendorf tube to await the next block. Each assay block required approximately 1–1.5 hours per batch so the total time individuals spent inside the Eppendorf tubes ranged from a few minutes to roughly 2 h, depending on their randomized order within each block.

This batch design was chosen to avoid individual marking methods (e.g., color-coding, ArUco, or QR codes), which would have posed logistical and biological challenges for a sample of 240 ants. The large arenas made HD video resolution inadequate for reliably detecting micro-tags or subtle color patterns from above, and physical marking requires anesthesia or delicate manipulation that risks injury, introduces acclimation delays, and prolongs isolation from the mother colony. To preserve individual tractability without marking, ants scheduled for the second day of each colony’s trial were collected before the first day’s subjects were returned to the nest. On completing all assays, individuals were either returned to the mother colony (if its testing was complete) or kept temporarily separate until the next batch was ready. This rotation prevented mixing of tested and untested ants, ensured all individuals were reunited with their colony within 24 hours, and avoided asocial rejection.

At the end of days 3 and 4, we took 4K resolution individual pictures of all ants that participated in the experiments. For each individual, we took measures of their head width (1.176 ± 0.082 mm), femur length (2.305 ± 0.377 mm), thorax length (2.154 ± 0.204 mm), and the length of the second segment of the antenna (1.644 ± 0.191 mm).

### 2.5 Data processing

Ants were tracked using ATracker [50], a Python toolkit designed for efficient, interactive tracking and processing of large batches of video files. Briefly, we applied automatic background subtraction and defined regions of interest and masks corresponding to the sociability chamber, the odor cue area, and the sucrose source. Then, batch tracking was performed to obtain x and y coordinates of the centroid of each ant at 10 fps. All videos were subsequently screened to confirm consistent detection of individuals, after which the data were automatically processed to extract, smooth, and convert trajectories, and to interpolate short gaps where ants were temporarily undetected. This ensured continuous and reliable trajectories for downstream analyses. We then calculated baseline instantaneous variables (speed, acceleration, turning angle, and distance to regions of interest) across all tracks.

To capture assay-specific spatiotemporal dynamics, we computed 24 additional metrics. Specifically, for the Exploration assay, we captured general locomotion by computing the average speed, distance traveled, proportion of time inactive, average direction persistence (mean cosine of consecutive turning angles), total meandering (sum of absolute turning angles divided by total path length), and reorientation rate (number of turns with angle > 150° by unit time). Spatial preference and coverage were measured via site fidelity to the arena center (area under the curve of the probability of finding the ant at a particular distance from a target location, in this case the center of the arena, measured across a range of distances), median distance to the center, the number of distinct sites visited (number of cells visited in a 5 × 5 mm grid), and the mean squared displacement (the average of the squared distance between the individual’s position and the center of the arena, across time windows of increasing length). For the Sociability assay, we quantified social affinity and group dynamics using site fidelity to the social chamber, average distance to the social chamber, the proportion of time interacting with nest-mates (a contact with the social chamber was considered an interaction), average interaction duration, and whether the tested individual exhibited at least one long interaction (continuous contact with the social chamber for more than 10 seconds). For the Chemical cue assay, we specifically evaluated sensory trail-following, using the proportion of time the ant was in the cue zone and its average distance to the edge of the zone across the trial. To analyze the preference to stay near the chemical cue, we measured the average distance to the cue and the proportion of time spent on the trail. To analyze the behavioral response, we calculated the average speed and reorientation rates both on the trail and in its vicinity (within 5 cm of the chemical trail, not including the trail itself). Finally, for the sucrose assay, foraging motivation and feeding persistence were assessed by measuring the proportion of time in the sucrose zone (the petri dish where the soaked cotton was placed), average distance to the nearest edge of the sucrose zone, and the latency to leave the sucrose zone. In total, we computed 24 metrics across the four assays.

### 2.6 Data analysis

To test whether ants differed according to their tasks in each assay, we fitted generalized linear mixed-effects models using `lme4` [51] and `glmmTMB`[52, 53]. We used average individual metrics as response variables and included task group and colony as fixed effects. The error structure was chosen according to the distribution of each response: Gaussian models for continuous metrics, logistic models for binary outcomes, and beta regressions for proportions. To account for the varying time individuals spent in the Eppendorf tube before testing, we included this duration as a continuous fixed effect; it was occasionally significant but always order(s) of magnitude smaller than the other predictors, and did not qualitatively alter the results, so we do not discuss it further (full model outputs in Table X). To account for variation associated with the experimental arena and time of testing during the day, random intercepts were included for setup and session.

Model assumptions were evaluated using the `DHARMa` package [54]. When assumptions were violated, models were refitted using transformed response variables (square-root, log, or second- and fourth-degree polynomial transformations). If this did not resolve the issue, non-parametric tests (Mann–Whitney or Wilcoxon rank-sum) were used instead, indicated in the results where relevant. The significance of fixed effects was assessed using likelihood-ratio tests. Estimated marginal means were obtained using `emmeans` [55], and post hoc contrasts were used to test specific hypotheses. When multiple comparisons were performed, p-values were adjusted using the Benjamini–Hochberg procedure for exploratory analyses, and the Tukey procedure for post hoc contrasts. To quantify feeding persistence, we used a log-normal accelerated-failure-time (AFT) model to assess how covariates accelerate or decelerate the time to departure from the food source and Cox proportional-hazards models to estimate the instantaneous probability of departure at any given time using `survival`, `survminer`, and `coxme` packages in R [56, 57].

Because even monomorphic workers can vary in size, we tested whether morphological variation could account for the behavioral differences we attributed to task. We summarized the morphological measurements into a single size axis using the first principal component of a PCA on the standardized traits, which captured X % of the morphological variance. We then asked, first, whether this size axis differed among task groups (using a single mixed-effects model with task group as a fixed effect and colony as a random effect), and second, whether including it as a covariate in the behavioral models altered the estimated task effects. Size did not differ significantly among task groups (Table Y), and its inclusion did not qualitatively influence the task effects on behavior, indicating that the behavioral structure we report is not driven by residual size variation.

The analyses above test whether task groups differ on individual behavioral metrics, one at a time. To instead ask whether individuals’ full multivariate profiles were structured by task, we built a behavioral landscape from our multivariate dataset (with the 22 behavioral metrics described above) using Uniform Manifold Approximation and Projection (UMAP) [58], a dimensionality reduction method that projects high-dimensional data into a low-dimensional space while preserving local similarity structure. This class of approaches allows complex, multivariate configurations to emerge organically from the data without relying on *a priori* categorization [35, 36]. To construct the embedding, the behavioral metrics were first centered and scaled, and subsequently, a principal components analysis (PCA) was performed to reduce dimensionality and noise. The first 15 principal components (explaining over 95% of the variance) were used as input for the UMAP embedding, computed with a neighboring parameter of 15 nearest neighbors. Discretization and clustering of the embedding were performed using a density-based, watershed-like algorithm implemented in the `bigMap` R package [59], which partitions the space according to local point density. The resulting clusters were then iteratively aggregated by merging smaller clusters with their nearest, higher-density neighbors, yielding four broad clusters that capture the main structure of the behavioral landscape. To quantify the distribution of behavioral profiles across tasks, we calculated the relative frequency of individuals from each task within each cluster. We tested whether groups differed in their distribution among the clusters using a global Chi-square test, followed by post-hoc evaluation of Bonferroni-corrected standardized residuals (|*z*| *>* 2.96) to identify specific cell-wise deviations from independence. Finally, to summarize and interpret the behavioral features of each cluster, we constructed separate PCAs for each assay using their respective behavioral variables. Scores from these assay-specific PCAs were projected onto the UMAP embedding and visualized as continuous color gradients, representing low to high levels of each behavioral dimension (see Results). All data analyses were conducted in R 4.4.2.

## 3 Results

### 3.1 Exploration

The trajectory data revealed a wide diversity of movement patterns, ranging from individuals spending long periods in the center of the arena to pronounced wall-following behavior and varying frequency of revisiting previously explored locations (Fig. 2A). This behavioral heterogeneity was reflected in the ants’ general movement metrics, with high inter-individual variability in average speed, turning rate, and stopping behavior within all ant groups (Fig. 2B). Ants from different task groups did not differ significantly in most movement parameters, with much greater within-group than between-group variability. Only average cruising speed (*χ*^2^ = 43.96, df =3, *P <* 0.001), and total distance travelled (*χ*^2^ = 15.56, df =3, *P* = 0.001) differed significantly among task groups, mainly driven by nurses moving at higher average cruising speeds than ants from other activity spaces (scouts *t*_21.8_ = 5.85, *P <* 0.001; recruits *t*_21.8_ = 3.19, *P* = 0.020; necrophores *t*_21.5_ = 5.61, *P <* 0.001). Nurses also traveled further than scouts (*t*_18.4_ = 3.47, *P* = 0.013) and necrophores (*t*_18.3_ = 3.13, *P* = 0.026), but not recruits (*t*_18.4_ = 2.34, *P* = 0.123). This heightened activity of nurses likely reflects a stress response to the abrupt transition from the nest to a bare arena, rather than a true exploratory behavior.

**Figure 2:**
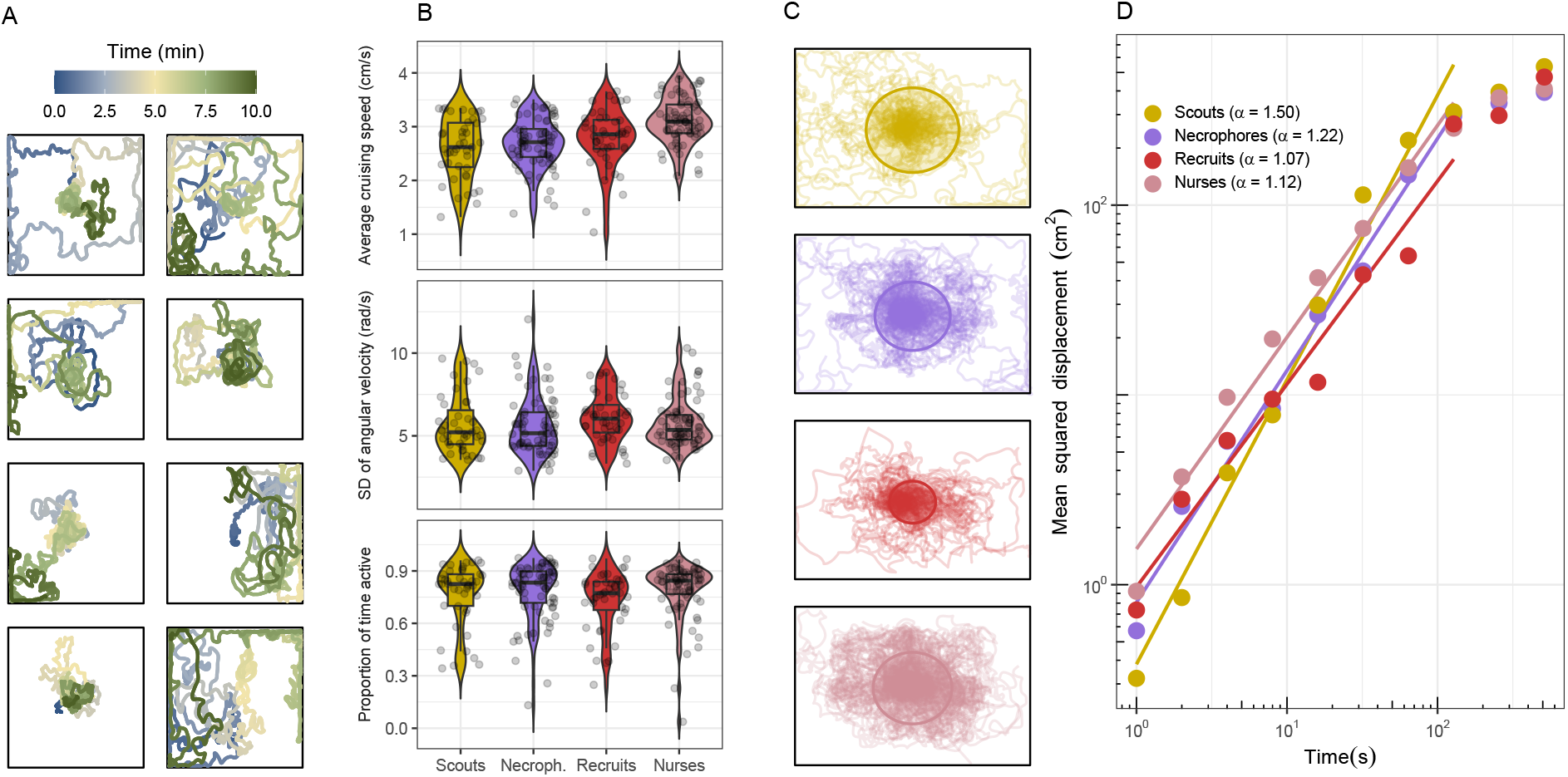
Characterization of exploratory activity in *A. senilis* workers. (A) Example trajectories illustrating a range of macroscopic exploratory patterns, from wall-following behavior (movement along the arena walls) to center-dwelling behavior (extended residence in the arena center). Color gradients indicate the temporal progression of each trajectory. (B) Subset of movement parameters used to quantify exploratory behavior in the open assays: average speed during movement (top), standard deviation of angular velocity (characterizing variability in turning behavior) (middle), and proportion of time spent inactive (bottom). (C) Trajectories of all individuals from each activity space (shown in the same colors and order as in D). The circle indicates the mean squared displacement reached at 2^6^ seconds (about 1 minute). (D) Mean squared displacement (MSD) computed over equally spaced time lags ranging from 2^0^ to 2^9^ seconds. The scaling exponents (*α*) were estimated from the MSD up to 2^7^ seconds (about 2 minutes), prior to the onset of border effects visible as saturation in the curves. Colors in B-D identify ants from the different activity spaces.

Two minutes into the trial, some scouts and necrophores reached the arena boundary, spreading out more widely than recruits or nurses did in the same period (Fig. 2C). Initially localized around the center of the arena, trajectories progressively expanded outward, a pattern effectively modeled as a diffusion process. The rate of this population-level expansion is captured by the Mean Squared Displacement (MSD, Fig. 2D), where the anomalous scaling exponent (*α*) classifies the movement regime as subdiffusive (*α <* 1), strictly diffusive (*α* = 1), or superdiffusive (*α >* 1) over time. Scouts (*α* = 1.50) and necrophores (*α* = 1.22) exhibited clear superdiffusive dynamics, demonstrating highly directional or persistent trajectories that accelerated outward-spreading over longer temporal scales. Conversely, recruits displayed a standard diffusive regime (*α* = 1.07), indicative of temporally unstructured random walks. While nurses occupied an intermediate position between diffusive and mildly superdiffusive regimes (*α* = 1.12), they remarkably yielded the highest diffusion coefficients (i.e., intercepts). This indicates that while their turning behavior resembles random walks, their high baseline speeds and displacement grant them highly localized coverage early on. Crucially, because *α* represents a power-law exponent, even small fractional increases in this value exponentially amplify long-term spatial spreading patterns as time progresses.

### 3.2 Social attraction to conspecifics

Ants exhibited substantial variation in the number and duration of visits to the social chamber, and their degree of engagement, with some individuals sustaining interactions for over ten seconds with their nest-mates. It was clear that overall, most individuals had a marked preference for remaining close to the social chamber, with a clear peak between 0 and 10 cm and a second peak up to 20cm away (Fig. 3A). Despite this spatial proximity, more than 50% of individuals interacted for less than 30 seconds over the course of the trial (Fig. 3B), with no differences in average interaction time among groups (*χ*^2^ = 2.79, df = 3, *P* = 0.424, Fig. 3B). Both the presence of heavy tails in the distribution of distance to nestmates (Fig. 3A) and the high density around 0 in the proportion of time interacting (Fig. 3B) suggested the presence of non-social individuals. These non-social individuals (i.e., ants that never approached within 1 cm of the social chamber at any point during the trial), were not evenly distributed among groups (*χ*^2^ = 8.42, *df* = 3, *P* = 0.038), with recruits showing the lowest proportion of non-social individuals compared to all other groups (Z ratio = 3.14, *P* = 0.002), with only two out of 44 individuals failing to interact (Fig. 3C). Together, the sociability metrics reveal a broad behavioral overlap across all task groups, but within it scouts to be the least social group and recruits the most social group, with necrophores more closely resembling scouts, and nurses more closely resembling recruits.

**Figure 3:**
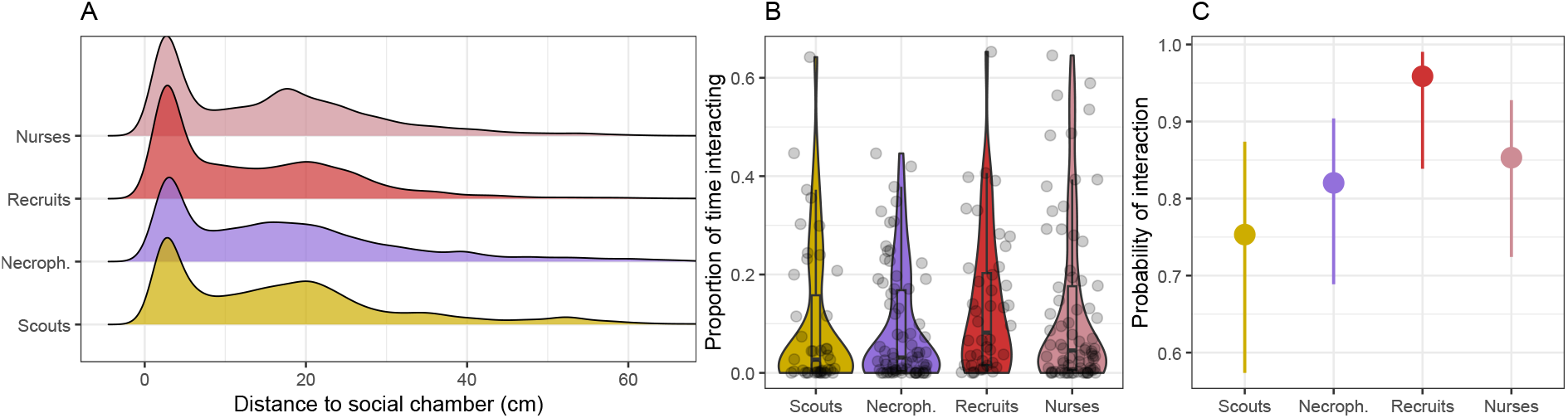
Characterization of sociability across individuals and activity spaces. (A) Distribution of distances to the social chamber, combining all distances recorded throughout the trial for all individuals across the different activity spaces. The densities are normalized by group so that the curves are comparable. (B) Distribution of the proportion of time spent interacting, i.e., within 1 cm (approximately one body length) of the social chamber. (C) Probability of interaction by group. Each dot represents an individual, classified according to whether they interacted with the social chamber (were found within 1 cm of the social chamber at least once in the trial) or not. The bars signify 95% CI of the estimated mean.

### 3.3 Responses to chemical cues

Chemical cues prompted a characteristic response in ants, with individuals typically following the odor gradient in a relatively straight trajectory and returning to the trail when the signal was lost. We found no significant differences among task groups in overall attraction to the cue, whether measured as the average distance to the cue (*χ*^2^ = 4.52, *df* = 3, *P* = 0.210; Fig. 4A) or the proportion of time spent on the trail (all Wilcoxon pairwise comparisons *P >* 0.05; Fig. 4B). The overlap in these distributions across groups suggests that individuals with both high and low responsiveness to the odor cue were present in all task groups. Nonetheless, nurses showed a slight tendency to remain closer to the trail, as reflected in a higher median proportion of time spent on it (Fig. 4A) and a greater concentration of positions near the cue, particularly within 0–10 cm (Fig. 4B).

**Figure 4:**
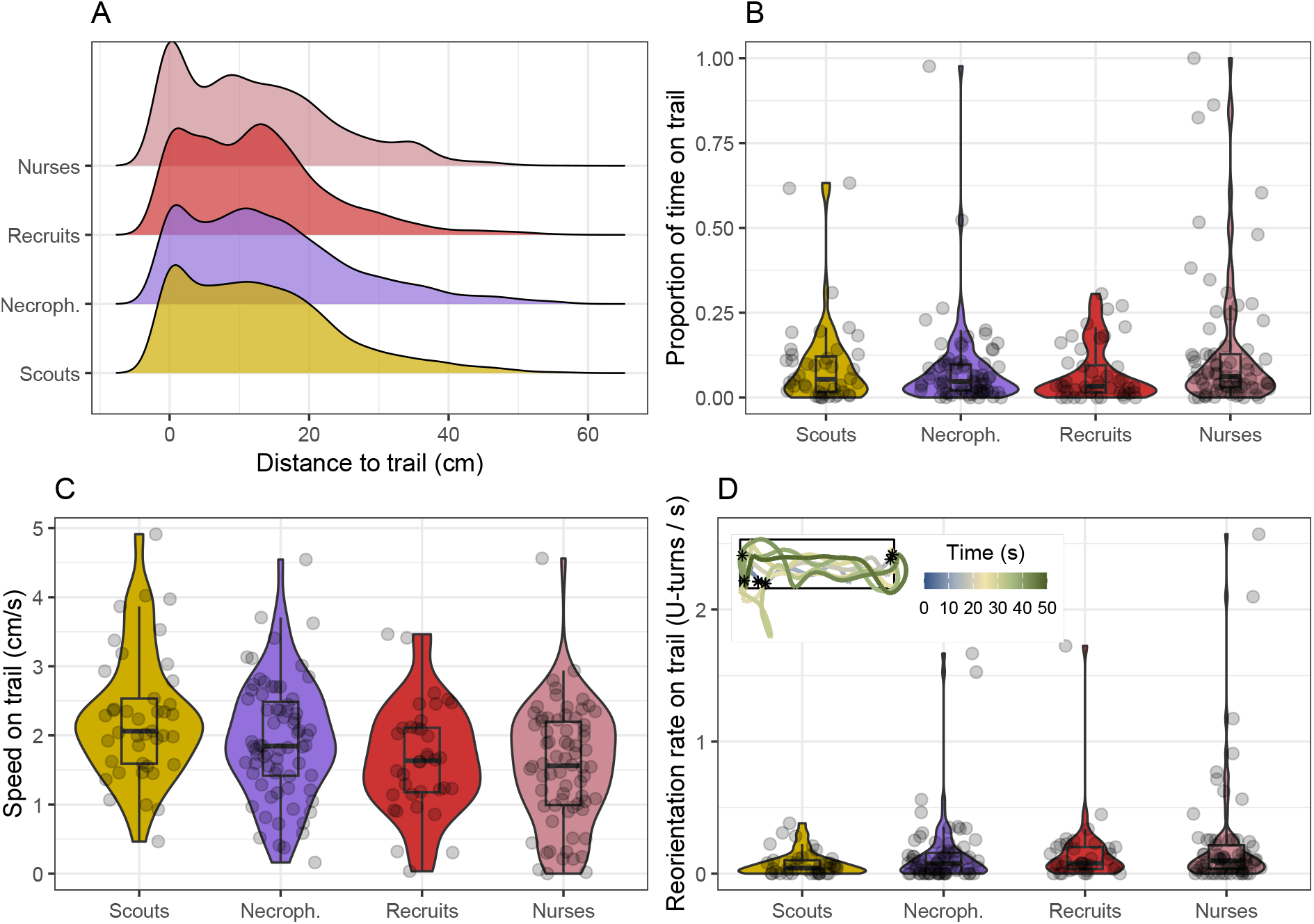
Characterization of attraction and reactivity to the odor cue across the four task groups. Distribution of distances to the odor cue, combining all distances recorded throughout the trial for all individuals across the different activity spaces. Densities are normalized within each task group to allow direct comparison. (B) Distribution of the proportion of time spent on the odor trail by each individual. (C) Distribution of individual average speed on the odor trail. (D) Distribution of individual sharp reorientation rates (U-turns). The inset displays an example track showing the characteristic trail-following behavior. The rectangle signifies the filter paper coated with the odor, and the stars show the spatial locations where a sharp reorientation occurred.

In contrast to the average responses to the cue, the locomotor reactivity differed significantly across task groups, as evidenced by differences in average speed on the trail (*χ*^2^ = 19.91, *df* = 3, *P* < 0.001; Fig. 4C). Recruits and nurses tended to reduce their speed in the presence of the cue, suggesting more careful signal tracking, potentially to refine the perceived direction of the gradient. This interpretation is supported by the spatial structure of the response (Suppl. Fig. 1): deceleration was most pronounced near the trail edges, where signal strength weakens. Across task groups, this deceleration was relatively mild in scouts and progressively stronger in necrophores, recruits, and nurses. In all cases, the reduction in speed was accompanied by an increase in turning rate near the cue (Fig. 4D). Although average turning rates did not differ significantly among groups (*χ*^2^ = 4.38, *df* = 3, *P* = 0.223), scouts consistently exhibited the lowest turning rates, while the remaining groups, particularly nurses, showed higher turning activity (Fig. 4D). Together, ants thus exhibited a gradient of preferences to remain on or near the cue trail that greatly overlapped among task groups, with cue-guided workers (recruits and nurses), as opposed to autonomous workers (scouts and necrophores), decelerating upon encountering or leaving the pheromone trail.

### 3.4 Responses to a liquid food source

Overall, there was substantial individual variation in the time ants spent feeding at the sucrose solution. For all task groups except for nurses, the distributions of feeding times were strongly bimodal, indicating a largely binary behavioral pattern in which individuals either showed little interest in the solution or remained at it for extended periods (Fig. 5A). This resulted in a significant difference among groups in the probability of feeding for more than 1 minute (*χ*^2^ = 28.50, df = 3, *P <* 0.001), with nurses exhibiting a lower probability compared with all other groups (Z ratio = -5.26, *P <* 0.001).

**Figure 5:**
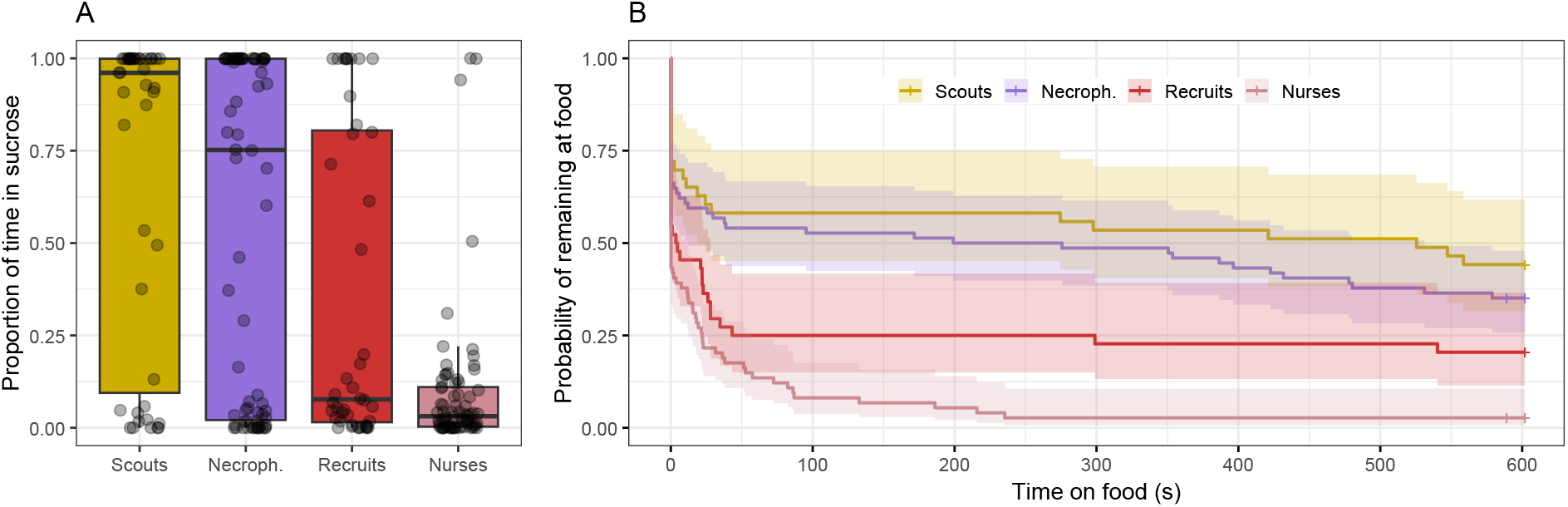
Propensity to feed by ants of different task groups. (A) Distribution of the time spent by individuals at the sucrose solution. (B) Survival curves depicting the probability of remaining at the food source over time. Mean estimate is shown as a line and the shades correspond to the 95% CI.

Similarly, survival curves showed that the probability of remaining at the food source over time varied markedly across task groups, with approximately 40% of scouts and necrophores remaining at the food for the entire duration of the experiment, compared with about 20% of recruits and only 2% of nurses (Fig. 5B). The log-normal accelerated-failure-time model revealed indeed that these differences were significant among groups (*χ*^2^ = 33.53, df = 3, *P <* 0.001). Relative to scouts, recruits exhibited substantially shorter feeding durations (time ratio = 0.054, *P* = 0.018), while nurses departed from the sucrose source even more rapidly (time ratio = 0.012, *P <* 0.001). These estimates indicate that recruits left the food source approximately 15 times faster than scouts, and nurses more than 85 times faster. Necrophores did not differ significantly from scouts (time ratio = 0.426, *P* = 0.358). These results were corroborated by the Cox proportional-hazards model (Suppl. Table 1), which showed that recruits were approximately twice as likely as scouts to leave the food source at any given time, while nurses were about times as likely. Feeding persistence and attraction to sucrose thus underscored a clear divergence in foraging priorities among groups, with scouts and necrophores displaying a strong attraction to the carbohydrate source, whereas recruits and particularly nurses largely disregarded the sucrose.

### 3.5 Behavioral profiles of *Aphaenogaster senilis*

We built a behavioral landscape that portrayed each individual as a unique behavioral profile. These profiles integrated individual metrics across the four experimental assays into a multivariate dataset. Clusters of varying densities emerged in the landscape, indicating a heterogeneous prevalence of the distinct behavioral profiles (Fig. 6). The landscape exhibited multiple organizational levels, with large isles formed by groups of clusters. For instance, clusters 2 and 6 formed one isle, while clusters 1, 7, 9, and 10 formed another, each containing finer-grained variability within.

**Figure 6:**
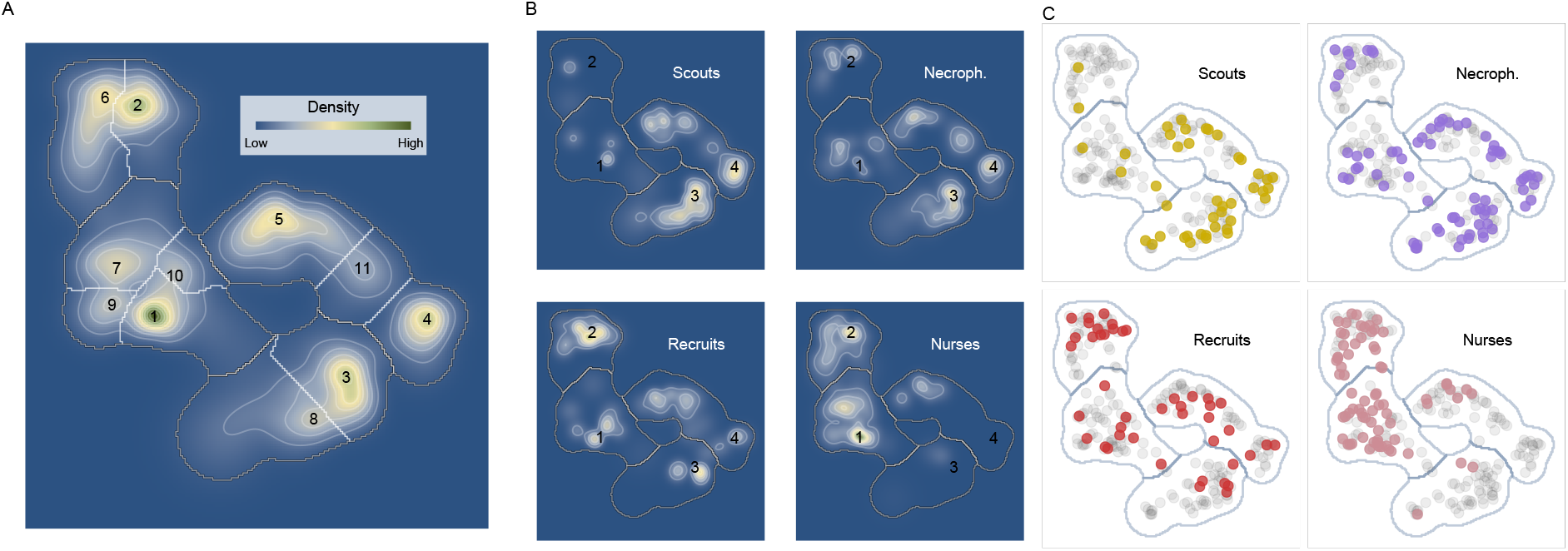
Behavioral landscape derived from UMAP. (A) Density heatmap of the embedding, ranging from low (blue) to high (green) density. Regions were segmented using a watershed algorithm (see Methods), yielding 11 clusters of varying density. The density peak of each cluster is labeled numerically according to its rank, and cluster boundaries are delineated by white lines. These clusters were further aggregated into four higher-level partitions (outlined with thicker dark gray lines) by iteratively merging clusters with their nearest, more dominant neighbors. The resulting partitions are labeled according to their dominant behavioral stereotype. (B) Density heatmap of individuals from each task across, normalized by the relative density of each task group. The numeric labels correspond to the merged clusters, and the lines outline their limits as in (A). (C) Raw data showing the distribution of individuals from each task across the behavioral landscape. Grey points indicate the full dataset, providing a reference for the overall embedding structure.

Mapping the task groups onto this behavioral landscape emphasized a spatial organization consistent with the patterns observed separately in each assay (Fig. 6B,C). Scouts and nurses were largely confined to specific clusters located in opposite regions: scouts predominantly occupied clusters 3 and 4 on the right, while nurses were concentrated in clusters 1 and 2 on the left. In contrast, recruits and necrophores were more sparsely distributed across the landscape.

To quantitatively assess the relationship between task groups and behavioral profiles, we calculated the relative prevalence of each task group within the clusters (Table **??**). A chi-square test rejected the null hypothesis of a homogeneous distribution of task groups across clusters (*χ*^2^ = 72.99, df = 9, *P <* 0.001), confirming significant differences in task distribution. Standardized residuals (Table **??**) showed scouts were under-represented in clusters dominated by nurses (clusters 1 and 2), and vice versa. Necrophores exhibited a similar, though less pronounced, pattern to scouts, mainly occupying clusters 3 and 4. Recruits were the only group whose distribution did not differ significantly from uniformity (*χ*^2^ = 4.00, df = 3, *P* = 0.261), being mostly evenly spread across all clusters.

We next identified the behavioral features defining each merged cluster, encompassing the four axes tested: exploration, sociability, and responsiveness to odor cues and sucrose (Fig.7, Suppl. Tab. 2, Suppl Fig. 2). Exploratory activity mainly decreased from left to right across the landscape, inversely mirroring the sharp increase in sucrose response along the same horizontal axis. For example, cluster 4 was characterized by low exploration but strong sucrose response, a trait it shared with cluster 3. Cluster 2 consisted of highly social individuals; however, social affinity measures, including sociability and odor cue responses, were dispersed throughout the landscape, complicating the interpretation of their spatial organization. These results indicate that exploration and sucrose-related traits, associated with foraging, are key discriminants of task groups, while sociability and odor cue responses, linked to communication, reflect individual variability rather than task specialization.

Because nurses, scouts, and, to a lesser extent, necrophores were predominantly associated with specific clusters, we examined the behavioral features characterizing each task group and the behavioral tendencies these tasks may select for (Fig. 7B). Nurses typically showed above-average activity during exploration assays and below-average interest in the sucrose solution. Two main nurse subgroups emerged: one highly social (approximately 36% of nurses, Cluster 2) and another with below-average sociability (approximately 49% of nurses, Cluster 1). Both groups exhibited average responsiveness to the odor cue, with the more social group (Cluster 2) showing slightly higher responsiveness. Scouts and necrophores were consistently characterized by above-average interest in sucrose. Within these groups, two classes also appeared: one with higher exploratory activity but lower sociability and odor cue responsiveness (about 42% of scouts and 34% of necrophores, Cluster 3), and another with average values across behavioral axes except for lower-than-average exploratory activity (about 42% of scouts and 35% of necrophores, Cluster 4). Note that around 30% of necrophores exhibited nurse-like behavior, roughly evenly split between the social (cluster 2) and non-social (cluster 1) nurse subgroups. Recruits were evenly distributed across these four clusters, behaviorally overlapping with the other task groups to varying degrees, with a slight tendency toward nurse-like behaviors (55% in clusters 1 and 2 versus 45% in clusters 3 and 4).

**Figure 7:**
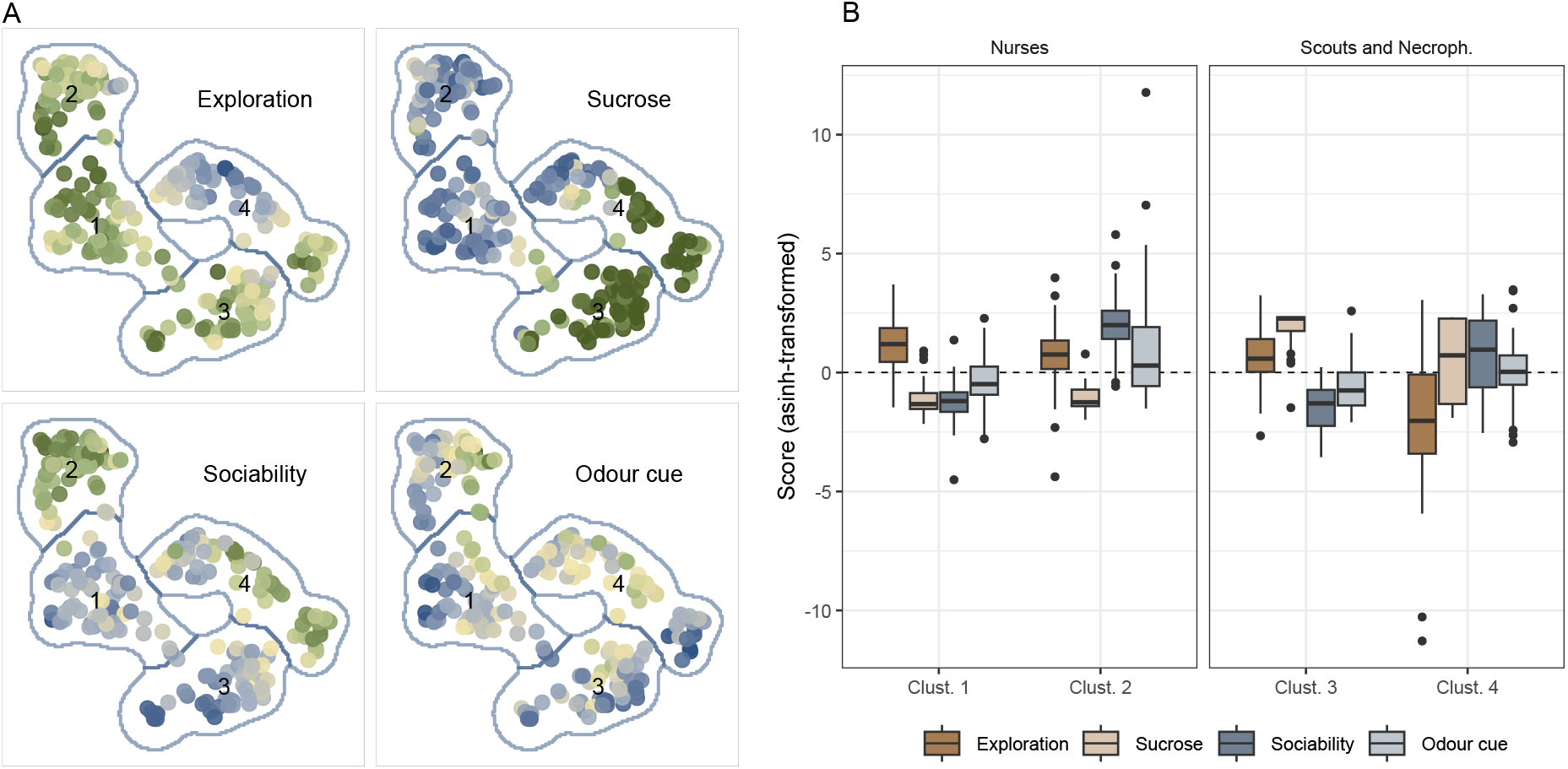
Behavioral profiles in the merged cluster partitions of the UMAP landscape. (A) UMAP embedding with individuals colored according to exploration, sociability, reactivity to odor cues, and sucrose scores. Each panel represents a linear combination of variables based on the loadings of the first principal component (PC1) from a PCA performed separately for each experimental assay (see Methods for details). Values were transformed using the inverse hyperbolic sine (asinh(*x*)) to reduce the influence of extreme values and enhance visual contrast. Values were then rescaled to the interval [−1, 1] within each assay to provide a common qualitative color scale, ranging from negative (blue) to positive (green). Distribution of these same variables across the merged clusters.

## 4 Discussion

Ants are often portrayed as highly stereotyped workers organized into castes that perform specific tasks, yet eusocial insects reallocate tasks dynamically in response to perturbations and environmental change. We therefore tested whether behavioral variability is constrained within task groups or occurs independently of task allocation. Using spatial position to infer functional roles, an approach grounded on the concept of activity spaces, we evaluated the behavioral profiles of ants engaged in four tasks (scouts, necrophores, recruits, and nurses) across ecological assays measuring exploration, sociability, and responsiveness to odor cues and sucrose. Our findings reveal a behavioral landscape lying between strict task specialization (division of labor) and complete behavioral flexibility (no association with tasks). Foraging-related traits, exploratory activity, and particularly sucrose responsiveness, were highly discriminative, cleanly distinguishing nurses from scouts, whereas communication-related traits (sociability and responsiveness to odor cues) varied substantially within task groups and overlapped broadly across them. Employing multivariate profiling and dimensionality reduction of individual behavioral profiles was key to uncovering the extent of this overlap and the organizational levels of behavioral variability. Overall, our results reveal behavioral constraints in scouts and nurses, greater flexibility in necrophores and particularly recruits, and set up the implications of these patterns for task transitions and colony resilience.

Our results support an intermediate view between strict task specialization and complete behavioral flexibility. On the one hand, we observed that behavioral heterogeneity was partially structured by task groups, with the most pronounced behavioral differences corresponding to extreme spatial positions deep inside (nurses) or far outside (scouts) the nest. On the other hand, we observed substantial behavioral heterogeneity within task groups, especially in the communication-based traits, resulting in considerable behavioral overlap across all task groups. Necrophores and particularly recruits were more sparsely distributed across the behavioral landscape, suggesting looser behavioral specialization in these groups. Arguably, this overlap could also arise if spatial sampling captured a mixture of task types at each location rather than genuine behavioral similarity. But the sharp, physiologically predicted sucrose gradient from nurses to scouts argues against this, indicating that the spatial assignment did capture real task differences.

High-levels of task specialization have been described in some social insect species, for instance, in preferences for collecting particular resources during foraging [60, 61, 46] or for tending larvae of a certain age [62]. This aligns with the classical perception of eusocial insects as being structured around stereotyped workers with clear preferences for performing certain tasks or roles, the so-called division of labor. However, theoretical and empirical work suggests that, in ants, task specialization scales with colony size and is limited or absent in most species [63, 18]. Even in species where specialization is strong and supported by morphological adaptations, individuals retain the ability to switch to unattended tasks when perturbations occur [64]. The idea of strong behavioral flexibility is also consistent with extensive findings showing low repeatability of behavioral assays in ants (e.g., [65, 66]). Importantly, such flexibility is not expected to reduce colony-level task efficiency [67, 32], and matches the foraging ecology of *A. senilis*: an opportunistic species that must rapidly exploit ephemeral resources before they are monopolized by dominant competitor species [30, 31].

Our findings sit between the two views: we detect task-structured differences, but against a backdrop of large within-group variation. Taken together, this evidence is more consistent with a framework in which task allocation emerges dynamically from spatially driven interaction networks than with the fixed behavioral specialization implied by classical division of labor [68]. Our results refine this view, however, by showing that the degree of behavioral specialization depends on the task and its spatial context. Crucially, this spatial context differs markedly across task groups. Scouts and nurses occupied the most distant regions of behavioral space, mirroring their occupation of the colony’s spatial extremes, while necrophores and recruits bridged these poles: necrophores more closely resembling scouts, and recruits more closely resembling nurses. This ordering parallels a gradient of environmental predictability, from the deterministic conditions of the nest interior to the stochastic world outside. The clearest structuring of behavior by task emerged along this gradient in foraging-related traits (i.e. sucrose and exploration assays) rather than communication ones (i.e. social and chemical assays). The behavioral divergence between scouts and nurses was accordingly most pronounced in foraging-related contexts. Nurses function within the nest under highly deterministic conditions, performing a constrained set of tasks associated with brood care [69, 46]. This environmental predictability plausibly explains their extreme and consistent responses in the foraging experiments: high exploration activity (likely reflecting an alert status and active search for brood-related cues) coupled with a lack of interest in sucrose, consistent with brood development relying primarily on protein rather than carbohydrate intake [46, 70]. In contrast, individuals operating closer to or outside the nest experience increasingly variable environments, reflected in a progressive increase in sucrose responsiveness from recruits to necrophores to scouts. This gradient in environmental stochasticity is mirrored in the spatiotemporal organization of movement, which shifts from diffusive to increasingly superdiffusive trajectories. Unlike diffusive movement, which tends to revisit already-explored areas, superdiffusive patterns combine local search with occasional long-range displacements, generating macroscopic patterns that improve encounter rates and exploration efficiency [8, 71, 72, 6].

In contrast to foraging-related traits, communication-related traits were largely similar across task groups, with within-task variability dominating over task-level differences. The only exceptions were that nurses slowed upon detecting chemical cues while scouts did not, and that recruits were slightly more likely to interact with nest-mates. Individual responsiveness to social information was otherwise highly variable within each task, with both socially responsive and non-responsive individuals present across all groups. This suggests that, while ants were broadly sensitive to social information, the propensity to engage in direct contacts was largely independent of task identity. Such variation has important consequences for colony-level connectivity: highly social individuals, regardless of task, increase the likelihood of inter-task interactions and act as bridges across functional groups, whereas less social individuals tend to accumulate contacts primarily within their own task group [40, 44, 45, 73]. This architecture naturally bridges short- and long-range connectivity [71], providing an additional mechanism through which eusocial insects may flexibly regulate information flow and tune collective responses to changing conditions.

A key methodological strength of this study is that behavior was quantified across large spatiotemporal scales, avoiding arbitrary categorical labels and capturing a richer, ecologically grounded behavioral space. Critically, the multivariate landscape revealed structure that the single assay analyses could not: whereas most individual metrics broadly overlapped across task groups, integrating them into whole-individual profiles exposed a clear task-related organization. This shows that the behavioral structure resides in the joint configuration of traits rather than in specific ones. This approach, based on variable coarse-graining, is moreover not limited to the task scale analyzed here: the same framework can navigate different organizational scales and recover behavioral structure at the individual, functional group, or colony level, offering a flexible tool for characterizing behavioral organization across colonies, species and contexts. Task groups were inferred from spatial position rather than direct behavioral observation, which carries an inherent limitation: we cannot be certain that spatial position always reflects the task an individual was performing at the time of sampling. However, this probabilistic approach was deliberately designed to minimize ambiguity, and as noted above, the sharp sucrose gradient from nurses to scouts supports the inference. Beyond this study, spatially-grounded sampling of this kind may offer a reproducible, non-invasive protocol applicable to other species and contexts.

Taken together, our results show that behavioural specialisation exists, but not across every dimension of behaviour, and it is not a fixed property of the task. These results suggest that workers may differ in their readiness to shift roles depending on the task at hand, and this variation may in turn shape how colonies adapt to environmental change. Broader behaviuoural profiles, associated with greater overlap between activity spaces, likely translate into greater ease of task switching. Within this framework, recruits and to a lesser extent necrophores would be expected to occupy transitional behavioral states that facilitate task switching, whereas workers engaged in more behaviorally constrained tasks, such as nursing or scouting, would be less likely to transition. This interpretation resonates with growing evidence showing that individuals occupying intermediate spatial positions act as key mediators of task transitions in social insects, functioning as behavioral stepping stones between more specialized roles [40, 44, 45, 73]. Our results extend this concept in *A. senilis*: given that necrophores are behaviorally scout-like and recruits nurse-like, we can hypothesize that necrophoring and scouting, on the one hand, and nursing and recruiting, on the other, are more readily interchangeable than other pairings of tasks. The learning costs associated with each task may also shape how interchangeable they are. The broad behavioral profile of recruits is particularly notable: they appear equipped to respond to a diverse range of stimuli, functionally resembling the reserve pools of inactive workers described in ant colonies [74, 23], which can be rapidly mobilized to cover unattended tasks. More broadly, maintaining a flexible workforce that moves across activity spaces and bridges spatially segregated groups may minimize colony-level task-switching costs, which would otherwise promote rigid specialization and likely reduce resilience to perturbation [75].

Our results speak to a question central to collective-behaviour research well beyond ants: how individual variability translates into functional structure at the group level. In this context, we suggest replacing the dichotomy of rigid behavioural stereotypes versus fully flexible workers with a richer, task-associated behavioural landscape. Individual heterogeneity, we show, is task-modulated, but with two important qualifications. First, the strength of this modulation depends on the task itself. Second, it does not extend to every behavioural dimension: highly social and non-social individuals coexist within all task groups, for instance, with the former likely acting as information bridges that facilitate communication both within and between tasks. Together, these findings point to a collective system that preserves stable group-level function while leaving room for flexible responses to emerge under specific ecological demands, a pattern likely relevant to any system in which adaptive collective function must arise from heterogeneous agents, from social insects to vertebrate groups.

## 5 Acknowledgements

This research has been supported by the Spanish government through Grant No. PID2021-122893NB-C21 (DO, FB and PFL).

