## Supplemental Figures and Tables for "Between specialization and flexibility: how tasks and space shape behavioral profiles in ant colonies"

July 20, 2026

### 1 Supplementary Tables

Table 1: Morphological measurements (mean ± SD) by behavioral group.

| Group | Head | Femur | Thorax | Antenna |
|---|---|---|---|---|
| Scouts | $1.199 \pm 0.073$ | $2.350 \pm 0.378$ | $2.176 \pm 0.236$ | $1.671 \pm 0.155$ |
| Necrophores | $1.174 \pm 0.081$ | $2.323 \pm 0.387$ | $2.131 \pm 0.213$ | $1.631 \pm 0.236$ |
| Recruits | $1.184 \pm 0.082$ | $2.277 \pm 0.401$ | $2.142 \pm 0.174$ | $1.652 \pm 0.173$ |
| Nurses | $1.155 \pm 0.090$ | $2.280 \pm 0.360$ | $2.174 \pm 0.199$ | $1.630 \pm 0.181$ |

Table 2: Pearson correlations ($r$) between morphological traits and exploratory behavior metrics. Bold values indicate statistically significant adjusted $p$-values ($P_{adj} < 0.05$), using a Benjamini-Hochberg protocol.

| Trait | Metric | $r$ | $P$ | $P_{adj}$ |
|---|---|---|---|---|
| Head | Average speed (cm/s) | 0.041 | 0.673 | 0.865 |
| | Proportion of time inactive | 0.036 | 0.709 | 0.865 |
| | Distance traveled (cm) | -0.002 | 0.984 | 0.984 |
| | Average directional persistence ($\cos(\alpha)$) | -0.233 | 0.014 | 0.164 |
| | Meandering (rad/cm) | 0.122 | 0.201 | 0.666 |
| | Reorientation rate (U-turns/s) | 0.116 | 0.226 | 0.666 |
| | Site fidelity (arena center) | 0.363 | $< 0.001$ | **0.002** |
| | Median distance to center (mm) | -0.359 | $< 0.001$ | **0.002** |
| | Number of visited sites (5×5 mm grid) | -0.172 | 0.071 | 0.513 |
| Antenna | Average speed (cm/s) | -0.099 | 0.296 | 0.666 |
| | Proportion of time inactive | 0.054 | 0.566 | 0.865 |
| | Distance traveled (cm) | -0.111 | 0.242 | 0.666 |
| | Average directional persistence ($\cos(\alpha)$) | -0.114 | 0.226 | 0.666 |
| | Meandering (rad/cm) | 0.022 | 0.820 | 0.923 |
| | Reorientation rate (U-turns/s) | 0.099 | 0.294 | 0.666 |
| | Site fidelity (arena center) | 0.161 | 0.088 | 0.526 |
| | Median distance to center (mm) | -0.178 | 0.058 | 0.513 |
| | Number of visited sites (5×5 mm grid) | -0.141 | 0.136 | 0.666 |
| Thorax | Average speed (cm/s) | 0.099 | 0.293 | 0.666 |
| | Proportion of time inactive | -0.009 | 0.928 | 0.984 |
| | Distance traveled (cm) | 0.051 | 0.591 | 0.865 |
| | Average directional persistence ($\cos(\alpha)$) | -0.044 | 0.643 | 0.865 |
| | Meandering (rad/cm) | -0.003 | 0.978 | 0.984 |
| | Reorientation rate (U-turns/s) | 0.028 | 0.768 | 0.892 |
| | Site fidelity (arena center) | 0.047 | 0.621 | 0.865 |
| | Median distance to center (mm) | -0.074 | 0.433 | 0.861 |
| | Number of visited sites (5×5 mm grid) | -0.034 | 0.717 | 0.865 |
| Femur | Average speed (cm/s) | 0.120 | 0.207 | 0.666 |
| | Proportion of time inactive | -0.043 | 0.651 | 0.865 |
| | Distance traveled (cm) | 0.084 | 0.374 | 0.793 |
| | Average directional persistence ($\cos(\alpha)$) | -0.121 | 0.202 | 0.666 |
| | Meandering (rad/cm) | -0.034 | 0.720 | 0.865 |
| | Reorientation rate (U-turns/s) | 0.006 | 0.947 | 0.984 |
| | Site fidelity (arena center) | 0.064 | 0.498 | 0.865 |
| | Median distance to center (mm) | -0.071 | 0.454 | 0.861 |
| | Number of visited sites (5×5 mm grid) | 0.035 | 0.711 | 0.865 |

| Context | Behavioral metric (unit) | Description |
|---|---|---|
| *Exploration* | | |
| | Average speed (cm/s) | Mean instantaneous movement speed over the assay. |
| | Proportion of time inactive | Fraction of time spent below a movement threshold (i.e., immobile). |
| | Distance traveled (cm) | Total path length covered during the assay. |
| | Average directional persistence ($\cos(\alpha)$) | Mean cosine of turning angle between steps; higher values indicate straighter trajectories. |
| | Meandering (rad/cm) | Angular change per unit distance traveled; higher values indicate more tortuous movement. |
| | Reorientation rate (U-turns/s) | Frequency of sharp directional reversals per unit time. |
| | Site fidelity (arena center) | Proportion of time spent in the central region of the arena. |
| | Average distance to center (mm) | Mean Euclidean distance from the arena center. |
| | Number of visited sites ($5\times5$ mm grid) | Number of distinct spatial bins visited, reflecting exploration breadth. |
| *Sociability* | | |
| | Site fidelity (social chamber) | Proportion of time spent in the compartment containing conspecific cues. |
| | Average distance to social chamber (mm) | Mean distance to the social stimulus area. |
| | Proportion of time interacting | Fraction of time engaged in physical or close-range interactions with conspecifics. |
| | Average interaction time (s) | Mean duration of individual interaction events. |
| | Cooperative behavior | Binary indicator (1/0) of cooperative behavior, defined as sustained actions aimed at liberating trapped nest-mates (e.g., biting the social chamber) lasting more than 10 seconds. |
| *Responsiveness to social cue (pheromone)* | | |
| | Proportion of time in pheromone | Fraction of time spent on the pheromone-marked area. |
| | Average distance to pheromone (mm) | Mean distance to the pheromone source or trail. |
| | Average speed on-trail (mm/s) | Mean movement speed while directly on the pheromone trail. |
| | Average speed near trail (mm/s) | Mean movement speed in the vicinity of the pheromone trail. |
| | Reorientation rate on-trail (U-turns/s) | Frequency of directional reversals while on the trail. |
| | Reorientation rate near trail (U-turns/s) | Frequency of directional reversals near (but not on) the trail. |
| *Food (sucrose)* | | |
| | Proportion of time in sucrose | Fraction of time spent in contact with or within the sucrose zone. |
| | Average distance to sucrose (mm) | Mean distance to the sucrose source. |
| | Time to leave sucrose (s) | Duration from first contact with sucrose to departure from the sucrose area. |

Table 3: Behavioral variables used to quantify exploration, sociability, responsiveness to social cues, and food-related behavior.

| Activity Space | Hazard Ratio | 95% CI | P |
|---|---|---|---|
| Necrophores | 1.26 | (0.77 - 2.06) | 0.355 |
| Recruits | 2.11 | (1.25 - 3.56) | 0.005 |
| Nurses | 3.57 | (2.22 - 5.74) | $< 0.001$ |

Table 4: Cox proportional hazards model estimating differences among activity spaces in the rate of leaving the food, with Scouts used as the reference group. Hazard ratios greater than 1 indicate faster departure than Scouts. Confidence intervals (95% CI) overlapping 1 indicate no statistically significant difference. Individuals remaining on the food at the end of the experiment were treated as right-censored (179 departures out of 235 ants). The proportional hazards assumption was satisfied (global Schoenfeld test: $\chi^2 = 7.41$, df = 3, $P = 0.06$)

| | Scouts | Necrophores | Recruits | Nurses | Total |
|---|---|---|---|---|---|
| Cluster 1 | 5 (7.7%) | 14 (21.5%) | 10 (15.4%) | 36 (55.4%) | 65 (27.7%) |
| Cluster 2 | 2 (3.8%) | 9 (17.3%) | 14 (26.9%) | 27 (51.9%) | 52 (22.1%) |
| Cluster 3 | 18 (34.6%) | 25 (48.1%) | 6 (11.5%) | 3 (5.8%) | 52 (22.1%) |
| Cluster 4 | 18 (27.3%) | 26 (39.4%) | 14 (21.2%) | 8 (12.1%) | 66 (28.1%) |
| Total | 43 (18.3%) | 74 (31.5%) | 44 (18.7%) | 74 (31.5%) | 235 (100%) |

Table 5: Counts (with row-wise percentages) by activity space and merged cluster from the UMAP landscape (Figs. **??** and **??**). Row totals and column totals (marginal values) are provided, with percentages relative to the row (cells) and to the grand total (margins).

| | Scouts | Necrophores | Recruits | Nurses |
|---|---|---|---|---|
| Cluster 1 | -2.60 | -2.03 | -0.81 | **4.88** |
| Cluster 2 | **-3.05** | -2.50 | 1.72 | **3.59** |
| Cluster 3 | **3.45** | 2.92 | -1.51 | **-4.53** |
| Cluster 4 | 2.22 | 1.63 | 0.61 | **-3.99** |

Table 6: Standardized residuals from the Chi-square test by merged cluster and activity space. Statistically significant deviations from independence are indicated in bold using a strict Bonferroni-adjusted threshold ($|z| > 2.95$, adjusting $\alpha = 0.05$ for 16 cell comparisons). Positive values indicate a significant over-representation (observed counts are higher than expected by chance), while negative values indicate a significant under-representation (observed counts are lower than expected by chance).

| Context | Variable | PC1 loading |
|---|---|---|
| *Exploration* | | |
| | Average speed (cm/s) | 0.299 |
| | Proportion of time inactive | -0.403 |
| | Distance traveled (cm) | 0.401 |
| | Average directional persistence (cos($\alpha$)) | 0.302 |
| | Meandering (rad/cm) | -0.370 |
| | Reorientation rate (U-turns/s) | -0.355 |
| | Site fidelity (arena center) | -0.222 |
| | Average distance to center (mm) | 0.196 |
| | Number of visited sites (5×5 mm grid) | 0.382 |
| *Sociability* | | |
| | Site fidelity (social chamber) | 0.490 |
| | Average distance to social chamber (mm) | -0.493 |
| | Proportion of time interacting | 0.497 |
| | Average interaction time (s) | 0.293 |
| | Cooperative behavior | 0.430 |
| *Responsiveness to social cue (odor cue)* | | |
| | Proportion of time in pheromone | 0.249 |
| | Average distance to pheromone (mm) | -0.229 |
| | Average speed on-trail (mm/s) | -0.491 |
| | Reorientation rate on-trail (U-turns/s) | 0.451 |
| | Average speed near trail (mm/s) | -0.487 |
| | Reorientation rate near trail (U-turns/s) | 0.450 |
| *Food (sucrose)* | | |
| | Proportion of time in sucrose | 0.603 |
| | Average distance to sucrose (mm) | -0.552 |
| | Time to leave sucrose (s) | 0.576 |

Table 7: Loadings of the first principal component (PC1) for each PCA used to interpret the distribution of exploration, sociability, responsiveness to social cues and sugar in the UMAP landscape.

#### 2 Supplementary Figures

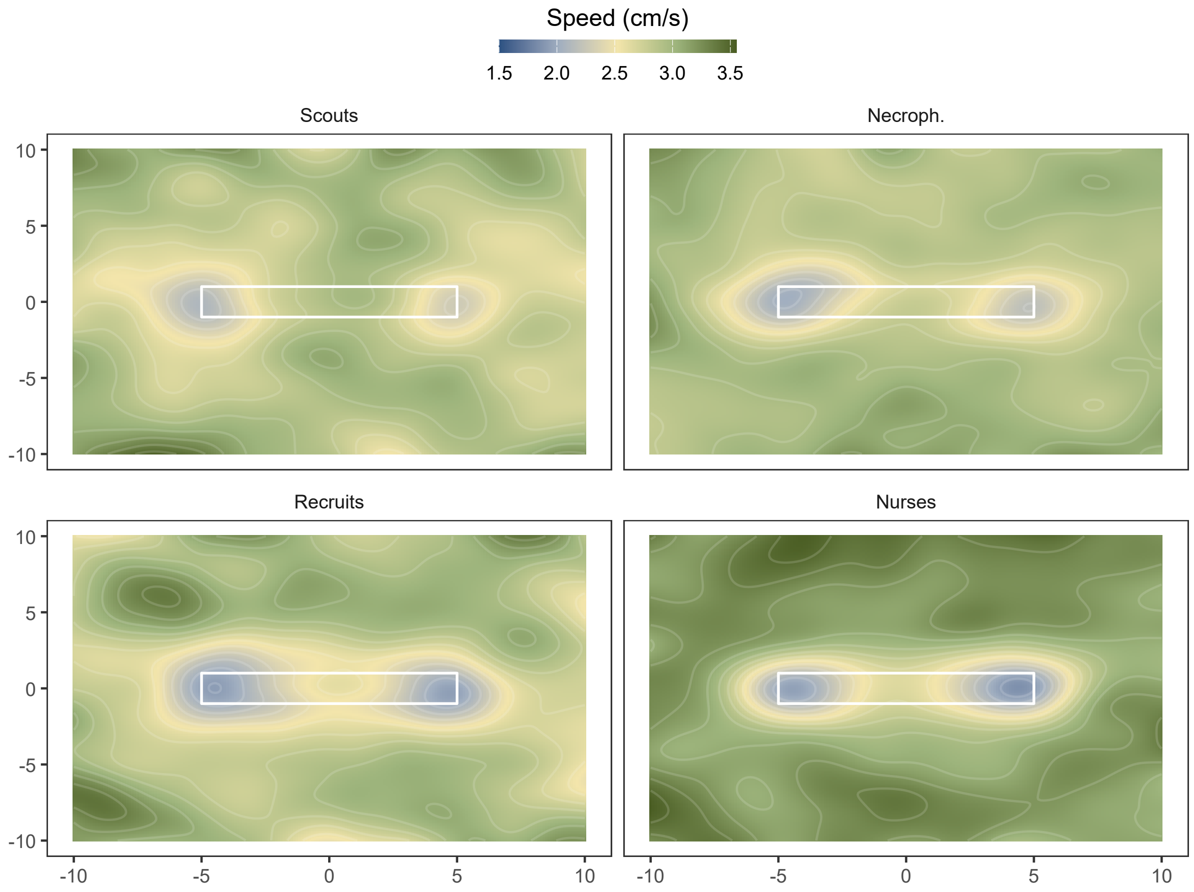


Figure 1: Heatmap showing the spatial distribution of speeds among groups in the odor cue assay. The white rectangle signifies the filter paper coated with the odor cue. The coordinates are in centimeters, and the origin of the coordinates is centered around the odor trail.

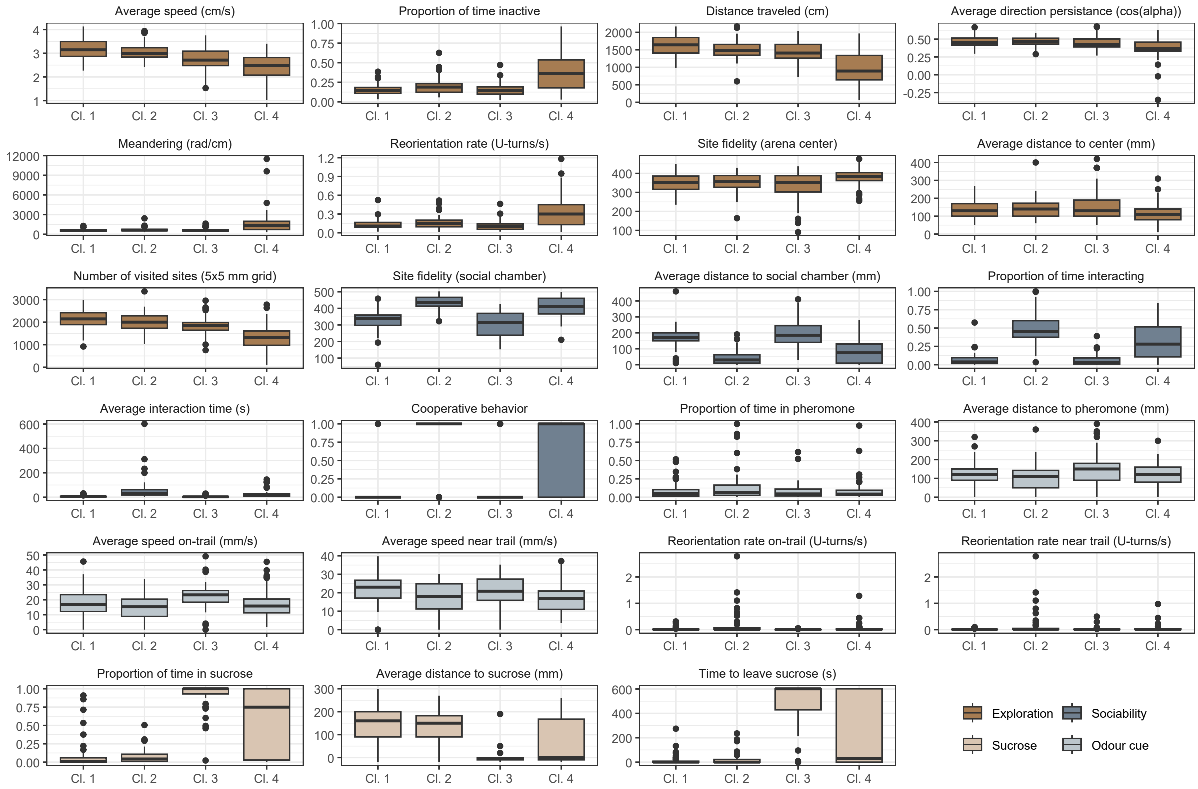


Figure 2: Distributions of the behavioral features used to construct the UMAP. The distribution of the variables are shown by behavioral cluster (as in Fig. ??). The variables are presented with units, and in the chronological order of the experimental assays: exploration, sociability, responsiveness to social cues, and food-related behavior. Boxplot colors indicate the experimental context (see legend).
